# Characteristics and data reporting of rare disease clinical trials: Getting better but still room for improvement

**DOI:** 10.1101/2021.12.03.471055

**Authors:** Nina K. Mair, Jürgen Gottowik, Raul Rodriguez-Esteban, Timothy J. Seabrook

## Abstract

**Background:** It is estimated that there are more than 7,000 rare diseases (RDs) worldwide, impacting the lives of approximately 400 million people and only 5% have an approved therapy. Facing special challenges, including patient scarceness, incomplete knowledge of the natural history and only few specialized clinical sites, clinical trials (CT) are limited, making the data from trials critical for research and clinical care. Despite the introduction of the U.S. Food and Drug Administration Amendment Act (FDAAA) in 2007 requiring certain CTs to post results on the registry ClinicalTrials.gov within 12 months following completion, compliance has been reportedly poor. Here, we describe general characteristics of RD CTs, identify trends, and evaluate result reporting practices under the FDAAA aiming to draw awareness to the problem of non-compliance.

**Methods:** CTs conducted between 2008 and 2015 were extracted from the public U.S. trial registry ClinicalTrials.gov using the text mining software I2E (Linguamatics). Disease names were matched with rare disease names from the Orphanet Rare Disease Ontology (ORDO, v2.5, Orphanet). Statistical analyses and data visualization were performed using GraphPad Prism 7 and R (v3.5). The Student’s t-test was employed to calculate significance using p-value cut-offs of <0.05 or <0.001.

**Results:** We analyzed 1,056 RD CTs of which 55.7% were phase 2, 7.7% phase 2/3 and 36.7% phase 3 trials. The studies were mostly one- and two-armed experimental CTs with the majority (60.2%) being funded by industry. Cystic fibrosis and sickle cell disease represented the most frequently investigated diseases (25.0% and 16.5%). Industry-led phase 2 RD CTs were significantly (p<0.0001) shorter than their equivalent led by academia/non-profit (22 vs. 33 months). Screening CTs completed before the end of 2015, we found that of the 725 analyzed studies, 55.2% predominantly phase 2 CTs, did not report results. Taking their potential applicability to the FDAAA into account, 25.2% industry-funded and 28.0% academia/non-profit-funded trials failed to disclose results on ClinicalTrial.gov.

**Conclusion:** RD CTs tend to be comparatively small, industry-funded studies focusing on genetic and neurologic conditions. Sponsor-related differences in study design, duration, and enrollment were observed. There are still substantial shortcomings when it comes to result publication.

## INTRODUCTION

In the U.S., rare diseases (RDs) are defined as those that affect fewer than 200,000 people; in Europe, the definition is based on a frequency of 1 in 2,000 or fewer people. It is estimated that there are more than 7,000 RDs which together affect the lives of approximately 400 million people worldwide^1^, and only half of these are genetically characterized^2^. The medical need is still high as only 5% of RDs currently have an approved therapy^3^. To promote research into these largely underappreciated orphan diseases, the U.S. Food and Drug Administration (FDA) introduced the Orphan Drug Act in 1983, which provides several incentives for the development of treatments^4^. Althoug this initiative has been very successful with 745 FDA approvals (428 in the last ten years), 99.9% of which have been submitted by industry, more therapies are needed^5^.

Clinical research in RDs faces challenges that typically have less impact on clinical trials (CTs) conducted in common diseases. These include patient scarceness, incomplete knowledge of the natural history of the disease, and the limited number of clinical sites that can treat these patients. This often results in a limited number of CTs in each RD, making the data from trials critical to research and clinical care. Despite the introduction of the U.S. Food and Drug Administration Amendment Act (FDAAA) in 2007^6^ requiring CTs matching specific criteria to post results on ClinicalTrials.gov within 12 months following trial completion, compliance has been reportedly poor. This may directly impact patient care and therapy development ^7–9^. The purpose of this report is to describe the overall landscape of rare disease clinical trials (RD CTs), identify trends, and assess the practice of result reporting after the implementation of the FDAAA to raise awareness of the problem of non-compliance.

## METHODS

### Data sources

Trial records with a clinical trial identifier (NCT number), a brief/official title and a study start and end date between 2008 and 2015 were extracted from the publicly accessible U.S. trial registry ClinicalTrials.gov using the text mining software I2E from Linguamatics. The glossary of common site terms, which can be found on ClinicalTrials.gov^10^, was used to determine if a CT was experimental, interventional, used an active comparator and other terms used in this paper.

### Dataset development

In order to distinguish RD CTs from common disease CTs, ClinicalTrials.gov condition specifications were matched with RD names from the Orphanet Rare Disease Ontology (ORDO) version 2.5 provided by Orphanet. Trials studying (rare) malignancies and communicable diseases were excluded, as well as all studies in phases 1, 1/2 and 4, and those without a study phase. Oncology CTs were excluded as rare oncology diseases are often included amongst other more common cancers as part of basket trials and thus did not reflect a RD CT. Infectious disease CTs included many HIV and hepatitis studies, which are not rare in much of the world and thus were excluded. Phase 1 studies comprise healthy subjects, not reflecting RD patient populations.

Studies starting before 2008 were excluded, due to the introduction of the FDAAA in 2007, as well as studies with a study status other than “completed”. Finally, studies involving incorrectly entered information into the “condition” field of the registry or studies that involved poisoning or envenomation were manually removed. To examine result reporting, we generated a second dataset including studies with an indicated completion date beyond 2015, to provide the responsible party with more than two years for the study results to be analyzed and entered on ClinicalTrials.gov at the time of our analysis.

To determine study location, all study locations in the contacts and locations section of each study’s ClinicalTrials.gov record were examined and counted. A site was counted every time it was mentioned by a CT. Therefore, a single study site was included multiple times if it hosted more than one CT during the time period examined.

To determine the sponsor, we extracted the responsible party as indicated on ClinicalTrials.gov and manually allocated CTs to “industry” or “academia/non-profit” according to the source of funding. “Industry” referred almost exclusively to registered for-profit companies, while publicly funded entities, such as universities, foundations, associations and non-profit organizations were allocated to “academia/non-profit”. One study in the final dataset of 1,056 studies could not be unambiguously allocated to a sponsor and was therefore excluded from the respective analyses.

### Statistical analyses and data visualization

Statistical analyses and data visualization were performed using GraphPad Prism 7. Microsoft Excel was used for statistical calculations to analyze enrollment due to cell number limitations in GraphPad Prism 7. For calculations of significance, Student’s t-test was employed using p-value cut-offs of <0.05 or <0.001 to demonstrate significant results.

## RESULTS

### Design characteristics of rare disease clinical trials (RD CTs)

An overview of the workflow resulting in the final dataset comprising 1,056 RD CTs is shown in **Fig 1**. Examining the general characteristics of the studies (**Table 1**), we found that trials were equally distributed across all years from 2008 to 2015 (12.5 ± 1.8% trials per More than half of the studies analyzed were in phase 2 (55.7%), followed phase 3 (36.7%) and 2/3 (7.7%). While phase 2 and phase 3 CTs showed a similar proportion of experimental (77.9% and 75.5%) and active comparator (17.3% and 21.7%) study arm types, phase 2/3 CTs tended to include more active comparator studies (29.6%) with 67.9% of the studies being experimental. Here, 49.4% of all CTs are two-armed studies, followed by one-armed studies (31.7%). 60.2% CTs were industry-funded and 39.8% were academia/non-profit-funded. While phase 2 and phase 3 CTs were predominantly industry-sponsored, with 53.9% and 74.1%, respectively, academia/non-profit prevailed in phase 2/3 CTs (60.5%).

**Figure 1.**
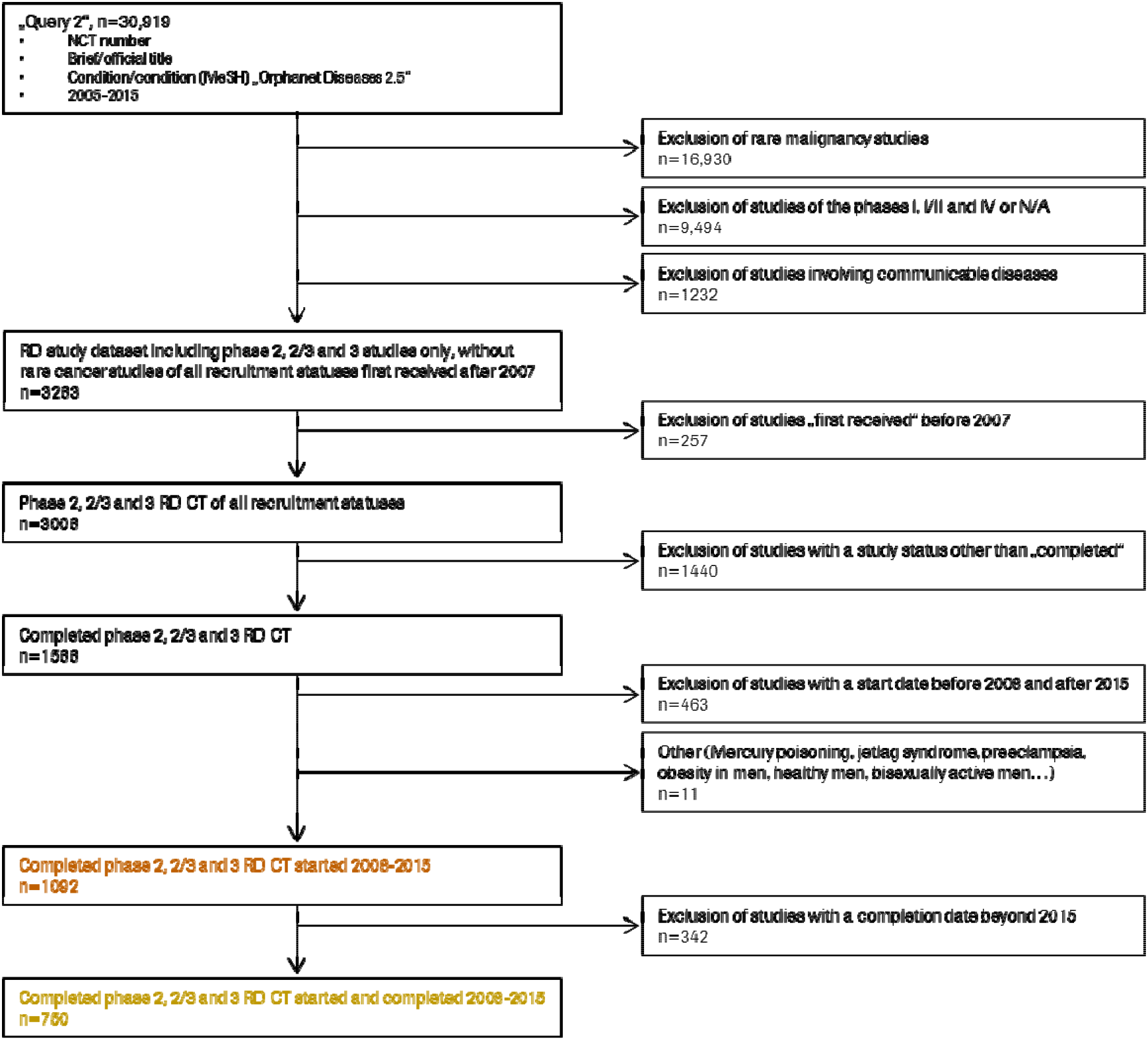
Flowchart showing the dataset development including filtering steps carried out to select the datasets for analysis.

**Table 1.** Characteristics of the 1056 completed Phase 2, 2/3 and 3 CTs registered at ClinicalTrials.gov. For each study arm type, the overall distribution of the evaluated CTs is highlighted in bold with the respective proportion of each study arm type listed below. The same applies to the table depicting the number of study arms.

| Study Start Year | n (%) | Study Phases | n (%) |
| --- | --- | --- | --- |
| 2008 | 149 (14.1) | Phase 2 | 588 (55.7) |
| 2009 | 138 (13.1) | Phase 2/3 | 81 (7.7) |
| 2010 | 137 (13.0) | Phase 3 | 387 (36.7) |
| 2011 | 146 (13.8) | **Number of Arms** | **n (%)** |
| 2012 | 144 (13.6) | **Phase 2** | **588 (55.7)** |
| 2013 | 130 (12.3) | 1 | 200 (18.9) |
| 2014 | 120 (11.4) | 2 | 256 (24.2) |
| 2015 | 92 (8.7) | 3 | 76 (7.2) |
| **Study Arm Type** | **n (%)** | 4 | 39 (3.7) |
| **Phase 2** | **588 (55.7)** | 5 | 10 (0.9) |
| Active Comparator | 102 (9.7) | 6 | 3 (0.3) |
| Experimental | 458 (43.4) | 8 | 1 (0.1) |
| No Intervention | 1 (0.1) | 10 | 1 (0.1) |
| Other | 14 (1.3) | 22 | 1 (0.1) |
| N/A | 13 (1.2) | N/A | 1 (0.1) |
| **Phase 2/Phase 3** | **81 (7.7)** | **Phase 2/3** | **81 (7.7)** |
| Active Comparator | 24 (2.3) | 1 | 19 (1.8) |
| Experimental | 55 (5.2) | 2 | 50 (4.7) |
| Other | 1 (0.1) | 3 | 7 (0.7) |
| Placebo Comparator | 1 (0.1) | 4 | 5 (0.5) |
| **Phase 3** | **387 (36.6)** | **Phase 3** | **387 (36.7)** |
| Active Comparator | 84 (8.0) | 1 | 116 (11.0) |
| Experimental | 292 (27.7) | 2 | 215 (20.4) |
| Other | 8 (0.8) | 3 | 37 (3.5) |
| Placebo Comparator | 1 (0.1) | 4 | 12 (1.1) |
| N/A | 2 (0.2) | 5 | 2 (0.2) |
|  |  | 6 | 4 (0.4) |
|  |  | 7 | 1 (0.1) |

### Disease categories

To generate a more comprehensive view of the RD CT landscape, we analyzed the disease areas addressed in this dataset. By leveraging the Orphanet Diseases 2.5 ontology, we were able to map all but one trial to specific conditions. Each CT could be allocated to up to seven diseases and each disease could be assigned to more than one category. The ten most common rare disease categories were genetic disease (19.4%), neurologic disease (12.3%), respiratory disease (8.2%), eye disease (6.4%), systemic or rheumatologic disease (5.9%), hematologic disease (5.4%), renal disease (5.0%), infertility (4.7%), skin disease (4.4%) and developmental defect (4.1%). These accounted for 75.8% of all diseases. In fact, 92.9% (n=971) of the RD CTs were mapped to at least one of the top 10 categories. In detail, 35.8% (n=378) of the studies were mapped to one top-ten category, 21.8% (n=230) were mapped to two and 20.5% (n=216) to three.

Furthermore, we found that the top ten diseases account for 86.2% of all CTs with cystic fibrosis (CF) accounting for 25.0%, followed by sickle cell disease (16.5%), hemophilia (8.7%), Fabry disease (6.3%), fragile X syndrome (6.1%), Huntington disease (5.7%), sarcoidosis (4.9%), systemic sclerosis (4.9%), Friedreich Ataxia (4.3%) and muscular dystrophies (primarily Duchenne, but also Becker and oculopharyngeal muscular dystrophy) (3.9%). To determine whether the ten most investigated diseases also reflect the leading diseases in each category, the various conditions were analyzed based on their by the sponsor designated category. While, CF dominates the “rare” categories genetic diseases (15.7%), respiratory diseases (37.1%) and infertility (64.2%), sickle cell disease was found first in the categories systemic or rheumatologic disease (16.9%) and renal disease (20.0%). The three most researched conditions in other areas are hemophilia (29.3%) in hematologic disease, alopecia (24.2%) in skin disease, and amyotrophic lateral sclerosis (12.1%) in neurologic disease.

Compared to the entire corpus of 2,257,370 CTs entered in ClinicalTrials.gov, most studies in ClinicalTrials.gov investigated cancers and other neoplasms (13.8%), followed by general pathology (10.5%), nervous system diseases (9.2%), digestive system diseases (6.7%), heart and blood diseases (6.6%) and behaviors and mental disorders (6.5%). Thus, RD CTs represent a different spectrum of disease areas as compared to more common diseases. In summary, the majority of the investigated diseases in our dataset are rare genetic and neurologic conditions with CF and sickle cell disease being the most studies diseases.

### Intervention type

Next, intervention types across study phases together with sponsors were analyzed (**Table 2**). Compared to ClinicalTrials.gov, where 45.6% (n=135,555) of all registered CTs involve drugs and biologicals as primary intervention type^11^, drug interventions (including small molecules) constituted 78.5% (n=829) of the CTs in our dataset, followed by biologicals (13.1%, n=138).

**Table 2.** Intervention Type per Phase and Sponsor for 1055 completed phase 2, 2/3 and 3 CTs registered at ClinicalTrials.gov. One study was removed from the analysis since the sponsor could not be assigned unambiguously to any of the two categories.

| Intervention Type<br>per phase | Industry |  |  | Academia/Non-Profit |  |  |
| --- | --- | --- | --- | --- | --- | --- |
|  | n (% per<br>Phase) | n (% per<br>Intervention Type) | n of CTs in<br>this phase (%) | n (% per<br>Phase) | n (% per<br>Intervention Type) | n of CTs in<br>this phase (%) |
| **Phase 2** | **316 (53.7)** |  |  | **272 (46.3)** |  |  |
| Behavioral | 0 (0.0) | 0 (0.0) | 0 (0.0) | 5 (0.9) | 5 (100.0) | 5 (1.8) |
| Biological | 45 (7.7) | 45 (73.8) | 45 (14.2) | 16 (2.7) | 16 (26.2) | 16 (5.9) |
| Combination Product | 1 (0.2) | 1 (100.0) | 1 (0.3) | 0 (0.0) | 0 (0.0) | 0 (0.0) |
| Device | 7 (1.2) | 7 (53.8) | 7 (2.2) | 6 (1.0) | 6 (46.2) | 6 (2.2) |
| Dietary Supplement | 1 (0.2) | 1 (11.1) | 1 (0.3) | 8 (1.4) | 8 (88.9) | 8 (2.9) |
| Drug | 257 (43.7) | 257 (54.6) | 257 (81.3) | 214 (36.4) | 214 (45.4) | 214 (78.7) |
| Genetic | 1 (0.2) | 1 (50.0) | 1 (0.3) | 1 (0.2) | 1 (50.0) | 1 (0.4) |
| Other | 1 (0.2) | 1 (11.1) | 1 (0.3) | 8 (1.4) | 8 (88.9) | 8 (2.9) |
| Procedure | 3 (0.5) | 3 (20.0) | 3 (0.9) | 12 (2.0) | 12 (80.0) | 12 (4.4) |
| Radiation | 0 (0.0) | 0 (0.0) | 0 (0.0) | 2 (0.3) | 2 (100.0) | 2 (0.7) |
| **Phase 2/Phase 3** | **32 (39.5)** |  |  | **49 (60.5)** |  |  |
| Behavioral | 0 (0.0) | 0 (0.0) | 0 (0.0) | 1 (1.2) | 1 (100.0) | 1 (2.0) |
| Biological | 14 (17.3) | 14 (87.5) | 14 (43.8) | 2 (2.5) | 2 (12.5) | 2 (4.1) |
| Device | 1 (1.2) | 1 (25.0) | 1 (3.1) | 3 (3.7) | 3 (75.0) | 3 (6.1) |
| Dietary Supplement | 0 (0.0) | 0 (0.0) | 0 (0.0) | 4 (4.9) | 4 (100.0) | 4 (8.2) |
| Drug | 16 (19.8) | 16 (30.2) | 16 (50.0) | 37 (45.7) | 37 (69.8) | 37 (75.5) |
| Other | 1 (1.2) | 1 (100.0) | 1 (3.1) | 0 (0.0) | 0 (0.0) | 0 (0.0) |
| Procedure | 0 (0.0) | 0 (0.0) | 0 (0.0) | 2 (2.5) | 2 (100.0) | 2 (4.1) |
| **Phase 3** | **287 (74.4)\*** |  |  | **99 (25.6)\*** |  |  |
| Behavioral | 0 (0.0) | 0 (0.0) | 0 (0.0) | 2 (0.5) | 2 (100.0) | 2 (2.0) |
| Biological | 60 (15.5) | 60 (98.4) | 60 (20.9) | 1 (0.3) | 1 (1.6) | 1 (1.0) |
| Device | 1 (0.3) | 1 (25.0) | 1 (0.3) | 3 (0.8) | 3 (75.0) | 3 (3.0) |
| Dietary Supplement | 1 (0.3) | 1 (16.7) | 1 (0.3) | 5 (1.3) | 5 (83.3) | 5 (5.1) |
| Drug | 225 (58.3) | 225 (74.0) | 225 (78.4) | 79 (20.5) | 79 (26.0) | 79 (79.8) |
| Other | 0 (0.0) | 0 (0.0) | 0 (0.0) | 5 (1.3) | 5 (100.0) | 5 (5.1) |
| Procedure | 0 (0.0) | 0 (0.0) | 0 (0.0) | 4 (1.0) | 4 (100.0) | 4 (4.0) |

In detail, 81.3% (n=257) of phase 2 RD CTs conducted by industry were found to involve drugs, followed by biologicals (14.2%, n=45) and devices (2.2%, n=7). Similarly, drugs and biologicals with 78.7% (n=214) and 5.9% (n=16) respectively, made up the majority of phase 2 CTs in the academia/non-profit sector. Notably, academia/non-profit-led trials tended to investigate intervention types beyond drugs or biologicals, such as dietary supplements, devices or procedures. Phase 3 RD CTs were found to be almost exclusively industry-sponsored and investigate drugs (78.4%) or biologicals (20.9%). Academia/non-profit CTs focused on drugs (79.8%), but not on biologicals (1.0%). In summary, the majority of studies investigate drugs with the main sponsor being industry, whereas academia/non-profit-led trials tend to explore a broader range of intervention types.

### Study location

The selection of investigational sites can be pivotal for enrollment and overall study success and their number varies considerably depending on the sponsor. Out of the 1,056 trials in our dataset, 632 (59.8%) entered at least one study site and, analyzing both sponsor categories separately, no significant difference was found. While 58.3% of all 635 CTs run by industry entered a study location, it was 62.1% of those run by academia/non-profit out of 420. One study could not be allocated to a sponsor and was therefore excluded leaving 1,055 trials for analysis. Although CTs from all around the world can be registered in ClinicalTrials.gov, we found that 44.3% (n=4,431) of the study sites were located in the U.S.. Here, the sum of counts by location does not equal the total CT number, as each location indicated by a study is counted and a single study may be counted more than once. Analyzing the number of sites by continent, the majority of sites are located in North America (48.5%, n=4,845) and Europe (33.9%, n=3,387), followed by Asia (11.4%, n=1,141), Central and South America (2.9%, n=286), Australia and Oceania (2.7%, n=269) and Africa (0.7%, n=72). The countries with the most study sites were the U.S. with 4,431 sites, Germany with 621 sites, France with 566 sites, Japan with 429 sites, Canada with 414 sites and the UK with 395 sites as well as Italy and Spain, with 326 and 251 sites, respectively. Approximately one third of RD CTs were run at a single site (34.7%, n=219), followed by 14.9% with up to five sites, 12.5% with up to ten sites and 9.7% with up to 15 sites. 4.6% and 5.4% of the studies had up to 15 and 25 study sites, respectively, and 1.6% indicated more than 100. The remaining 16.8% were scattered between 25 and 100 study sites. Moreover, industry-led trials significantly increased the number of study sites with ascending study phase. Accordingly, the mean location number for industry in phases 2, 2/3 and 3 was 15.0 (median of 7.5), 23.0 (median of 18.0) and 32.7 (median of 23.5), while for academia the location numbers per phase were 3.1 (median of 1.0), 4.6 (median of 1.0) and 9.6 (median of 1.0). Compared to the entire trial corpus present on ClinicalTrials.gov, where 40% of the CTs indicated trial sites in the U.S. (35% U.S. only and 5% U.S. and non-U.S.), we found that RD CTs tend to be more often conducted in the U.S. (44.3%). In summary, the vast majority of trials entered into ClinicalTrials.gov are conducted in North America and Europe and despite a correlation between a higher numbers of trial sites and industry funding, single-site trials are the most common.

### Participant enrollment

CT patient recruitment is critical and can be especially challenging in RD CTs due to patient scarceness and dispersion. The enrollment numbers in our dataset follow the typical trend observed in common diseases in that the number of participants increases with progressing study phase. Participant numbers were heterogeneous, as reflected by considerable standard deviation values. The median patient recruitment for industry CTs in phases 2, 2/3 and 3 was 40.0 (interquartile range (IQR), 21-76), 77.5 (IQR, 50-227), and 80.5 (IQR, 37-191), respectively, vs. 26.0 (IQR, 12-47), 35.0 (IQR, 20-88) and 60.0 (IQR, 23-110) for academia/non-profit CTs. This enrollment difference between sponsor types was significant (p<0.05) in phases 2 and 2/3. The mean enrollment was approximately 96% (industry) and 83% (academia/non-profit) higher than the median. Of the 73,071 participants enrolled in industry-funded trials, 59.4% had been recruited for phase 3 CTs, followed by 33.6% for phase 2 CTs, and 6.9% for phase 2/3 CTs. Academia/non-profit enrolled 44.0% of 24,959 participants in phase 3 CTs, 40.7% in phase 2 CTs, and 15.3% in phase 2/3 CTs. In analyzing gender eligibility of RD CTs, we found that the majority of the studies, 89.3%, admitted participants of any gender, whereas 7.1% and 3.6% of RD CTs enrolled only male and female subjects, respectively. In sum, industry-led trials consistently enrolled consistently more patients than academia/non-profit-funded trials, with notable variability as reflected by the standard deviation in enrollment.

### Participant Age

A RD can affect anyone irrespective of age, but at least half of RDs manifest themselves in early childhood. To determine if this is reflected in RD CTs, the participant age was assessed. Therefore, the entries for the ClinicalTrials.gov categories “participant age”, “participant minimum age” and “participant maximum age” were retrieved (**Table 3)**.

**Table 3.** Participant minimum and maximum age as indicated by RD CTs in our dataset (n=1056).

| Participant Minimum Age | n (%) | Participant Maximum Age | n (%) |
| --- | --- | --- | --- |
| < 1 year | 24 (2.5) | < 1 year | 15 (2.5) |
| 1-10 years | 204 (20.8) | 1-10 years | 31 (5.2) |
| 11-18 years | 637 (64.9) | 11-18 years | 90 (15.2) |
| 19-35 years | 66 (6.7) | 19-35 years | 47 (7.9) |
| 36-65 years | 50 (5.1) | 36-65 years | 164 (27.6) |
| 65< | 0 (0.0) | 65< | 247 (41.6) |
| **Total** | **981 (100)** | **Total** | **594 (100)** |

A minimum age for participants was entered for 92.9% (n=981). Of the CTs showing a minimum age, 88.2% allowed patients under 18 years to be enrolled. In contrast, only 56.3% of the studies indicated an upper age limit for participants. Of those, 41.6% indicated an extended age of eligibility above 65 years and a notable fraction (15.2%) of trials set the maximum age between 11 and 18 years. While extracting the minimum and maximum age limits was straightforward, the analysis of the participant age, filled out by 96.8% of the studies, proved complicated due to the lack of uniformity of the information entered. After manual categorization into age groups, we were able to visualize the data (**Fig. 2**). Even though around three quarters of the studies set the upper age limit to more than 60 years, numerous studies put their focus on a younger population, mirrored by the high proportion of studies indicating a minimum participant age below 18.

**Figure 2.**
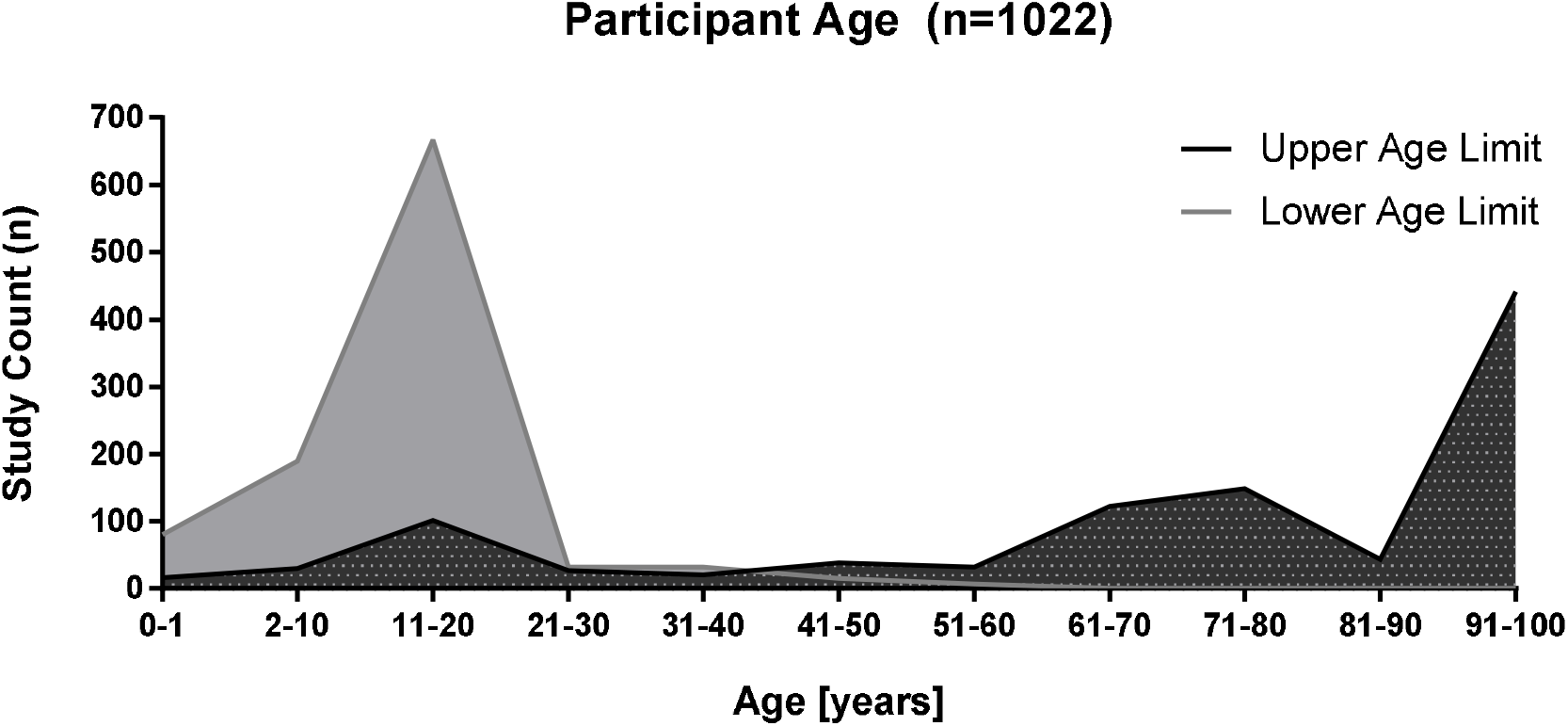
Graphic representation of age restrictions for participants of the RD CTs in our dataset. Of the total of 1,056 CTs that were analyzed, 1,022 entered age specifications.

### Study duration

Another aspect of CTs is study duration, as calculated in this analysis from the study start to the study completion date. We found that phase 2 studies conducted by industry were, with a median duration of 22 months (IQR, 14-33), significantly (p<0.0001) shorter than their academia/non-profit-led equivalent, with a median duration of 33 months (IQR, 22-47)(**Fig. 3**). This trend could be similarly observed in phase 3 CTs, with industry conducting shorter studies with a median of 27 months (IQR, 18-41) versus 34.5 months (IQR, 19-52) for academia/non-profit CTs (p<0.05). In contrast, the study duration did not differ significantly in phase 2/3 CTs between industry and academia/non-profit with a median of 29 (IQR, 21-37) and 31 (IQR, 18-48), respectively. In summary, industry-funded phase 2 and 3 RD CTs are of a shorter duration than studies led by academia/non-profit.

**Figure 3.**
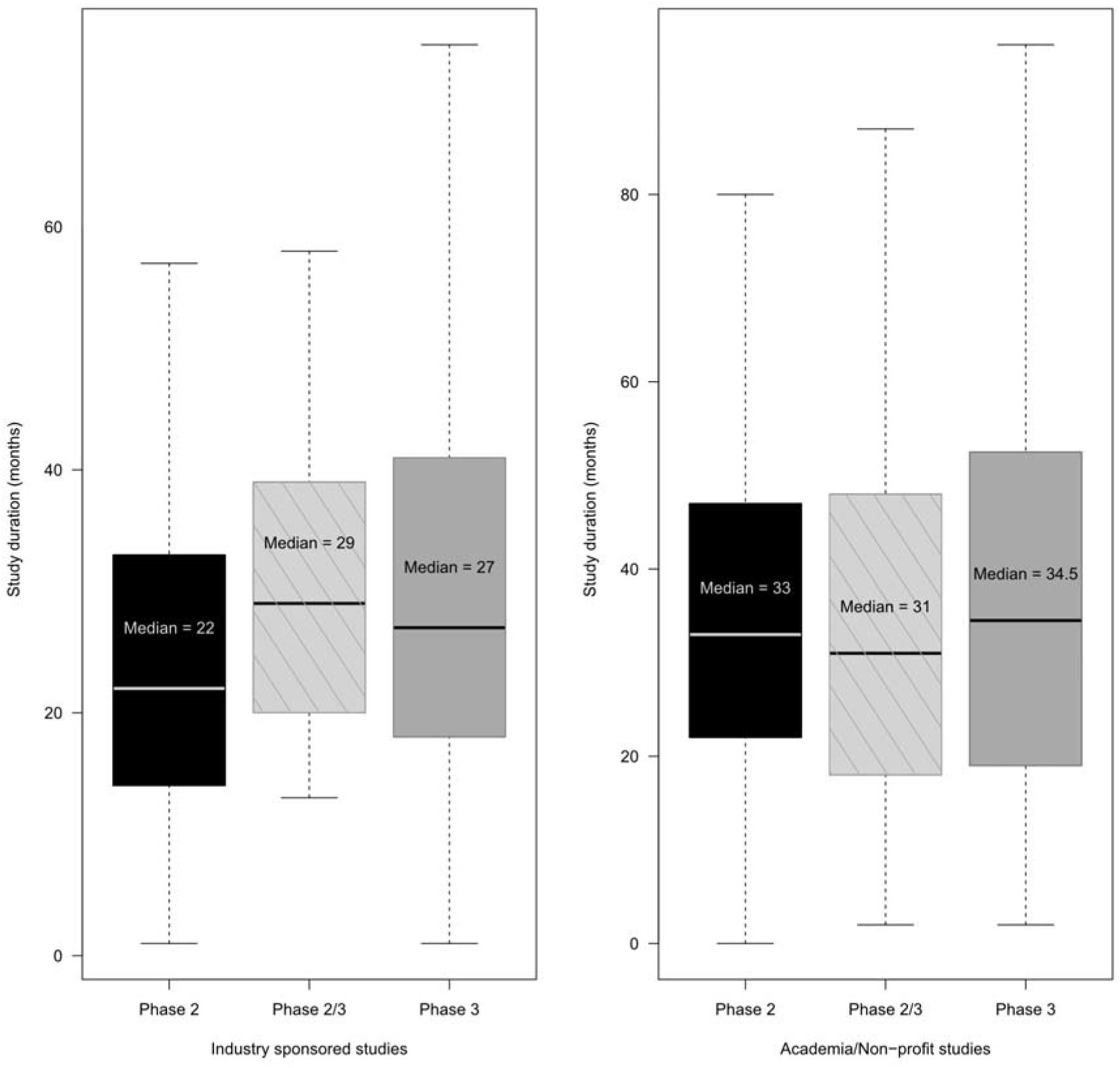
Comparison of study duration in months between industry- and academia/non-profit-sponsored RD CTs. This analysis is based on the dataset comprising 1,056 studies.

### Result reporting among rare disease clinical trials

In 2007, the FDAAA was introduced requiring the posting of results for eligible studies on ClinicalTrials.gov within twelve months after study completion (FDAAA section 801). A CT qualifies as eligible or applicable to the FDAAA if the drug, biological or device under investigation has been manufactured in the U.S. or its territories, or if the CT has at least one study site in the U.S. or its territories. To examine result reporting, a second dataset of (n=725 CTs) consisting of 55.9% phase 2, 7.6% phase 2/3, and 36.6% phase 3 CTs was generated (**Supplementary table 1**). Overall, 44.8% (n=325) trials had entered results while 55.2% (n=400) had not. Of those that failed to report results, 65.8% were phase 2, 9.8% were phase 2/3, and 24.5% were phase 3 CTs. Industry reported consistently more results over all years (2008-2015) with a total of 56.3% (n=243) reporting RD CTs. Strikingly, only 28.0% (n=82) of studies conducted by academia/non-profit did likewise. To explore if sponsors in the non-reporting dataset were meeting their obligations under the regulations, we extracted the RD CTs which were identified as including at least one U.S. site (n=134) or using a drug manufactured within the U.S. (n=1). For the remainder of the studies (n=265) determining if the FDAAA was applicable could not be easily concluded from the information included in ClinicalTrials.gov, thus a random sample of approximately 25% (n=67) were examined manually. The majority (n=53, 79.1%) of the studies in this sample were not applicable to the FDAAA, while four (6%) CTs were identified as applicable. The remaining ten (14.9%) studies in this sample did not provide enough information for a clear categorization. We found that one of the studies that was not applicable displayed the status “results submitted” with no results entered in the respective entry field meaning that the results might currently be under evaluation. Extrapolating the manually extracted dataset of 67 studies to the results of the overall number of non-reporting trials, we estimated that 20.8% of the included 725 CTs failed to report results although required by U.S. law. Thereof, 105 CTs (24.3%) are industry-funded and 46 CTs (15.7%) academia/non-profit-funded studies. Adding trials with unclear applicability (0.9% industry- and 12.3% academia/non-profit-led CTs), the overall proportion of result non-reporting CTs, although (potentially) legally required, increases to 25.2% and 28.0% for studies initiated by industry and academia/non-profit, respectively. In summary, more than half of the industry-funded and roughly one third of academia/non-profit-funded RD CTs report results online after study completion. Of those that unambiguously fall under the regulation of the FDAAA, CTs sponsored by academia/non-profit enter results more often than industry-led CTs, with the latter posting more results online on a voluntary basis. However, including also potentially applicable CTs, result reporting practices do not differ between sponsors.

## DISCUSSION

We conducted a landscape analysis of completed phase 2, 2/3 and 3 RD CTs registered in the ClinicalTrials.gov database, including CTs from all disease areas except oncology and infectious diseases. RD CTs were shown to be mostly sponsored by industry, which is in line with previous findings in more common diseases^12^. Academia/non-profit CTs prevail in phase 2/3 possibly because CTs in this phase often explore already approved drugs in new populations, or for extended indications. This falls in line with a previous observation of academia-led trials peaking predominantly before phase 3^13^. The present industry dominance in phase 3 CTs, which often are needed for regulatory approval, can be linked to those trials characteristically being large and of longer duration rendering them costly. Similar to previous reports, drug interventions represent the most frequent type of intervention in our dataset^14^. Unsurprisingly, industry focused on drugs and biologicals in all phases, with biologicals more common in phase 2/3. Academia/non-profit, although similarly focused on drugs, explored a broader array of interventions, including devices and dietary supplements. That might be partially due to the challenges posed by the nature and diversity of the dietary supplements branch and its globally varying regulations among different jurisdictions or governmental dominance in this field^15^. The strong presence of academia/non-profit sponsors in CTs around biologicals and drugs could be due to research done on approved drugs aiming to extend their scope towards new applications or patient groups in RDs. As previously reviewed^16^, pharmaceutical companies and academics are entering into collaborations to repurpose approved or discontinued drugs for other indications, including RDs. This may explain the increased numbers of non-industry sponsored phase 2/3 trials. Overall, comparing our data with similar data in ClinicalTrials.gov, no differences between our RD dataset and the CTs in more common diseases could be detected and intervention types employed in RD research, with the largest group being phase 2 CTs investigating drugs conducted by industry, reflect the common practice in general clinical research^17^.

Most RD CTs are genetic or neurologic diseases. The high prevalence of genetic CTs in the RD dataset is to be expected, as the majority of RDs are thought to be genetic in origin^18^. Among these, CF is the most researched disease, representing also the most common genetic disease in Caucasian children counting 70,000 cases worldwide. This substantial number of patients, paired with early screening policies and a molecular understanding of the disease facilitates the setup of CT and thereby therapy development^19–20^, leading pharmaceutical companies to conduct studies in this field^21^. Similar reasons might apply for neurologic disorders, the second focus of CTs in our dataset, which occur relatively frequently in the pediatric population^22^. These observations are in line with what can be found for common diseases present in ClinicalTrials.gov, where nervous system disorders are the most researched disease area, followed by neoplasms and general pathologies. In contrast, digestive system diseases, heart and blood diseases, and behaviors and mental disorders were highly represented in ClinicalTrials.gov, but not in our RD dataset. Therefore, although rare diseases are numerous, the vast majority of clinical trials focus on a few disease categories and conditions, such as CF. Furthermore, we found Huntington disease (HD) representing the top disease in rare eye disorders, followed by uveitis. Although HD is not primarily an eye disorder, a possible explanation for this finding could be a common endpoint in this disease (Unified Huntington’s Disease Rating Scale), including ocular assessments causing its misclassification as eye disorder. This example highlights the difficulties that can occur when analyzing data from Clinicaltrials.gov that has not been curated or reviewed critically.

In line with a recently published study^23^, we identified North America and Europe as the regions and the U.S. and Germany as the countries with the most study locations. The high proportion of U.S.-led CTs could be explained by the strong presence of the pharma industry compared to other countries with the ensuing availability of financial and infrastructural resources as well as the strong presence of active patient associations and foundations that play a key role in advancing research^21^. Japan, Australia, Argentina and South Africa were the countries with most CTs in Asia, the Oceania region, Middle and South America and Africa, respectively. However, 40.2% of CTs did not report a location, biasing the data and its interpretation. In accordance with a previous report analyzing the International Clinical Trial Registry Platform (ICTRP), we found Japan to be the country with the most CT sites in Asia^23^. While the mentioned report attributes the shift away from “traditional” CT sites in Western countries to lower study costs in other countries, the study location for RD CTs could be more influenced by patient prevalence, the availability of specialized care centers and specific legislation fostering RD research^24–25^.

Considering the general scarceness of patients and experts/specialized care centers in RDs, it is not surprising that half of the trials indicate at most five sites. In contrast, almost two thirds of the trials indicated more than one study site, pointing towards the implementation of more, but smaller study sites in RD research. Interventional RD CTs that were previously found to have less study sites compared to non-RDs might provide support for this hypothesis^26^. The correlation we found between industry-funded CTs and an increased number of study sites could be linked to the different financial support received compared to academia/non-profit. Although phase 2 CTs prevail in our dataset, most participants were enrolled in phase 3 CTs, with the majority enrolled in industry sponsored CTs. A study analyzing common disease CTs of all phases reported an overall enrollment rate below 100 participants for 62% of studies, which was the same percentage we observed in phase 3 CTs in ClinicalTrials.gov. Examining phase 2 and 2/3 RD CTs, the proportion of studies enrolling 100 or less patients rose to 87.8% and 72.8%, respectively. Comparing our dataset to the aforementioned study based on the respective median enrollment, we found comparable accrual numbers between RD CTs overall and therein described oncology trials, whilst mental health and cardiovascular CTs were higher at 85 and 100, respectively. It is somewhat surprising that the number of patients in RD CTs and non-RD CTs are similar, as we expected RD CTs to enroll fewer patients as previously indicated in a study comparing rare with non-rare disease CTs^26^. While this study describes mostly early phase CTs with fewer participants, most of the patients in our dataset were enrolled in phase 3 CTs. Additionally, the limited power of small studies to show significant clinical efficacy, might be avoided by study sponsors resulting in higher accrual numbers. Reasons for less participant accrual in non-RD CTs could be competing trials targeting similar participant populations or even an increased focus on small sub-populations within a larger disease area. This indicates that CTs in RDs and more common diseases may be becoming more similar and both may have issues with recruitment of sufficient numbers of patients to draw firm conclusions. Notably, we observed great heterogeneity between trials enrolling few patients, sometimes in single digits and trials enrolling hundreds or thousands, a phenomenon that has also been previously reported^17^.

Gender disparity in biomedical research, which may insert a bias in clinical findings and therefore may lead to a disadvantage in clinical practice for women, is recognized and has been described before ^17,27,28,29^. Although we did not observe a significant gender bias in RD CTs, we found that out of all studies recruiting only patients from a specific gender, twice as many recruited only male patients than female, which could be partially attributed to the aforementioned historic and general underrepresentation of women in CTs.

The age of eligibility for study subjects is crucial in many diseases, often influencing the overall study success. The majority of RDs are thought to be genetic disorders, which present already in early childhood or adolescence, thereby necessitating an early therapy start^30–32^. Consequently, RD CTs had a wide age range with the majority including pediatric patients, unlike in more common diseases that can occur throughout life^17,26^.

CT participation is often associated with great efforts by (pediatric) patients and their family members, thus the study duration can have a major impact on participant compliance and retention in the study. We found a direct correlation between the study duration and study phase, which is in line with common clinical research practices such as monitoring procedures, larger sample sizes, time to analyze larger datasets including side effects etc. in phase 3 studies. The fact that industry-funded CTs are significantly shorter than academia-/non-profit-led CTs could be attributed to the former usually having more resources, which allows for more sites, thereby increasing patient recruitment velocity. Comparing RD CTs to all phases of nephrology or cardiology studies, no significant difference could be detected in the overall study duration^14^.

Clinical research is essential for clinical decision making and providing the best standard of care for patients. Since the enactment of the FDAAA in 2007, CT result reporting is no longer only ethically desirable, but also mandatory within twelve months upon trial completion. However, this rule only applies to studies investigating a drug, biological or device manufactured in the U.S. or studies indicating at least one study site in the U.S. or within its territories^6^. Our analysis of studies completed before 2015 showed that after at least two years following completion less than half of all analyzed RD CTs posted results on ClinicalTrial.gov. A previous study on result reporting under FDAAA regulations found that only 22% of the CTs report results timely, with only 10% doing so on a voluntary basis. Additionally, the authors stated that result reporting occurs in only 40% of industry-funded and 9% not solely industry-funded CTs with phase 3 studies being the most frequently reported CTs^7^. Despite opposition of the FDA^33^, an unofficial analysis conducted by the U.S. National Institutes of Health (NIH) confirms that industry performs better than public sponsors (NIH and NIH-funded), when it comes to timely result reporting, with 52% vs. 35%, respectively^34^. A recent study analyzing the availability of phase 3 and 4 RD CT results on ClinicalTrial.gov found that 68% of the trials affected by the FDAAA reported results^35^. In our dataset, phase 2 trials account for roughly two thirds of the non-reporting CTs we identified. Studies registered with the EU Clinical Trials Register (EUCTR), which requires result disclosure within 12 months after trial completion, performed similarly, with only half of the applicable CTs posting results^36^. This recent report also found that CT result reporting was linked to later study phases and industry funding. Drawing a final conclusion on the result reporting rate between the two sponsor categories in our dataset is not obvious due to the incompleteness of the information provided online. Considering only studies that clearly apply to the FDAAA, industry funding is shown to be associated with a higher degree of non-reporting in RD CTs. However, including potentially applicable CTs, academia/non-profit and industry fall abreast. This importantly highlights the need for complete information in order for external parties to be able to draw transparent conclusions. Generally, industry trials in RDs were associated with better overall result posting even when not required by the FDAAA, potentially due to more strictly regulated and overseen follow-up processes and the fact that industry-funded trials tend to be larger and larger studies are more likely to be published^37^.

The finding that RD CTs adhere more strictly to the FDAAA than CTs overall might be due to ethical obligations towards these patients and the high value of clinical data from a CT, which may be the only one performed in a patient population. RD CTs also include many pediatric patients and pediatric CTs were found more likely to be completed^38^. Regardless of the reasons, it is encouraging to note that RD CTs are reported at a higher rate than other CTs. Ultimately, CTs need to overcome the widely recognized and general deficit of consistent and transparent data sharing practices, not only to improve patient care, but also to value the participants who help advancing science and future patients.

## CONCLUSION

This study provides insight into the landscape of RD CTs. RD clinical research tends to be industry-funded and focused on treatments with drugs or biologicals and many trials involve a relatively small number of participants and patients starting from very early in life. Generally, industry funding has been associated with larger studies, including larger numbers of participants and more study sites, but slightly shorter study duration compared with academia/non-profit trials. Finally, CT results in rare diseases are being made often publicly available more frequently than has been reported for studies entered in ClinicalTrial.gov, but improvements are still needed to make all data readily available to the public.

### Limitations

There are some limitations in this study to be taken into consideration. First, ClinicalTrials.gov does not cover all trials conducted worldwide. However, there is a reported 80% overlap between ClinicalTrials.gov and the WHO ICTRP portal^18^. Secondly, there is no single standard ontology for the description of clinical research and, despite extensive manual data curation efforts, the dataset may still contain misclassified studies. Conversely, some CTs may have been excluded from our study, due to the disease under study not conforming to the ORDO naming convention. Efforts were made to identify, correct or remove erroneous or ambiguous as well as carelessly entered data prior to analysis, however, an element of uncertainty remains. Additionally, a notable fraction of studies may provide divergent information on ClinicalTrials.gov and EUCTR, making comparisons difficult^39^. Although the FDAAA was introduced to increase transparency, this legislation came with certain limitations; For example, its applicability to studies with at least one study site or an item manufactured in the U.S. or U.S. territories. It is, however, not evident which studies are applicable and which ones are exempt. Even though we attempted to address this limitation by examining a random sample of 25% of the trials manually, errors may still be present. In this study, early proof-of-concept trials, which can be impactful on the scientific community, e.g. in order to avoid duplicity of resources, are not reviewed. Finally, this study represents a snapshot of RD CTs as they were entered in March 2018 and some study details may have been added after we retrieved the data.

## SUPPLEMENTARY MATERIAL

**Supplementary table 1.**
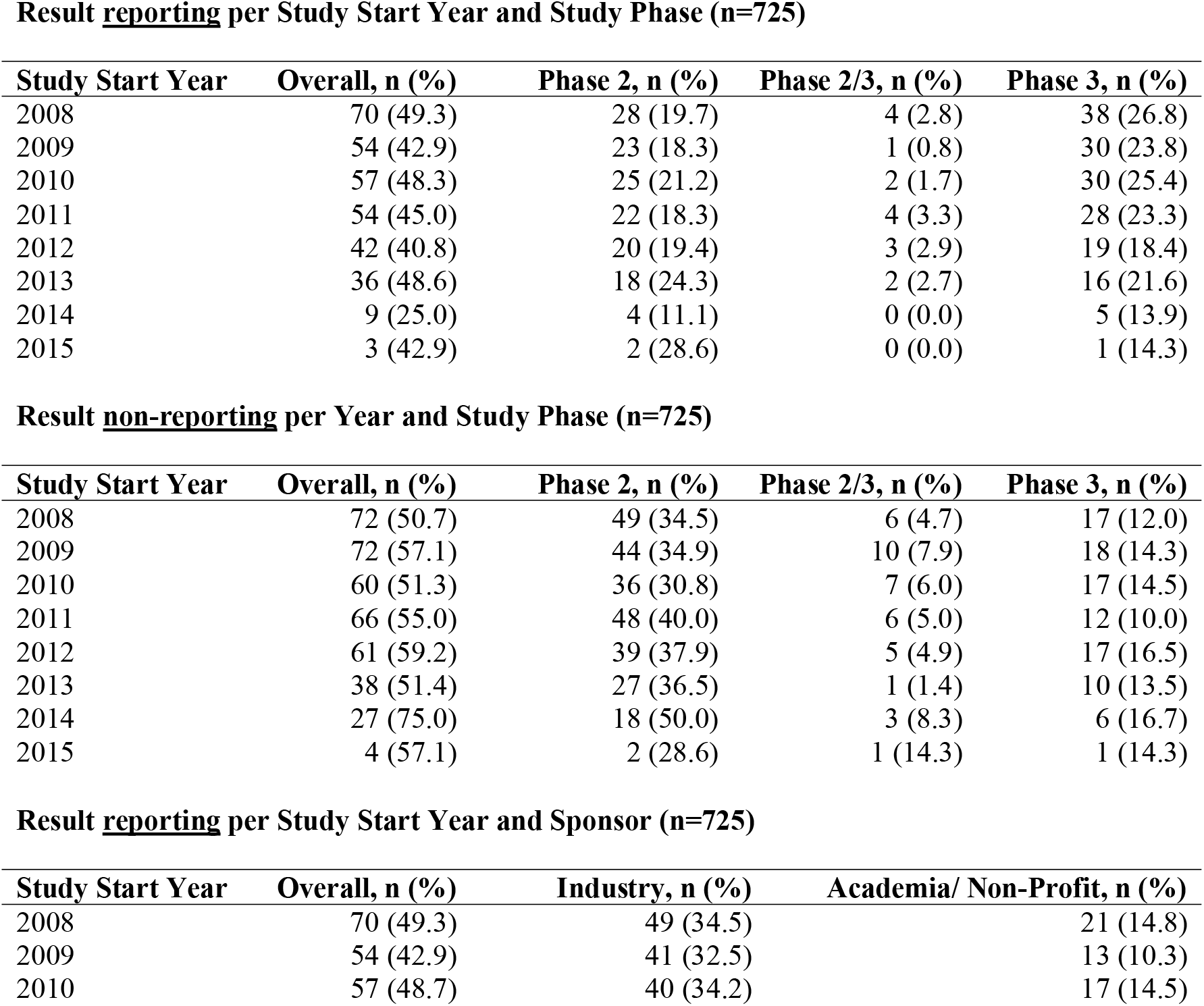
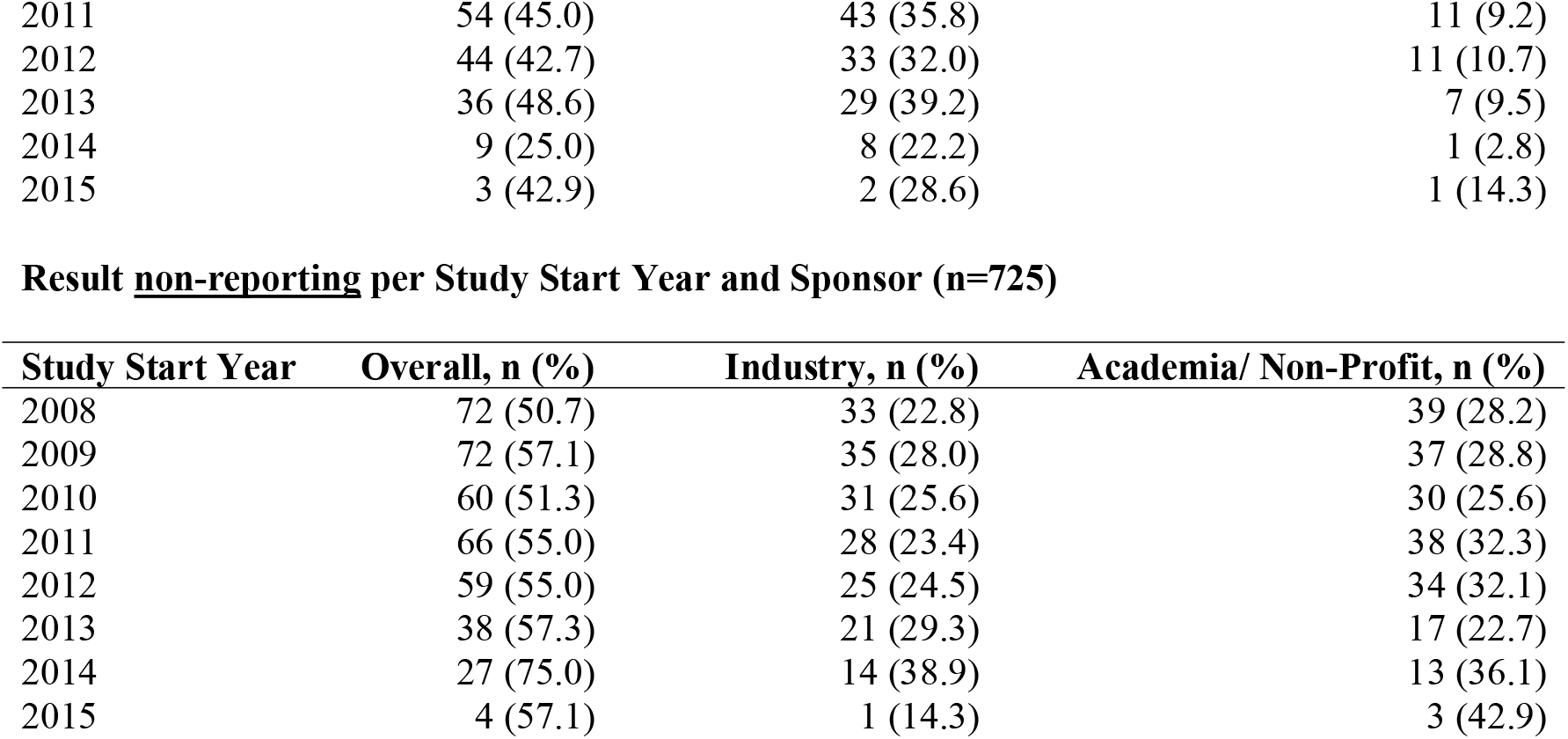
Clinical trial result publication/non-publication per study start year and sponsor irrespective the eligibility of the specific study.

Result reporting per Study Start Year and Study Phase (n=725)
| Study Start Year | Overall, n (%) | Phase 2, n (%) | Phase 2/3, n (%) | Phase 3, n (%) |
| --- | --- | --- | --- | --- |
| 2008 | 70 (49.3) | 28 (19.7) | 4 (2.8) | 38 (26.8) |
| 2009 | 54 (42.9) | 23 (18.3) | 1 (0.8) | 30 (23.8) |
| 2010 | 57 (48.3) | 25 (21.2) | 2 (1.7) | 30 (25.4) |
| 2011 | 54 (45.0) | 22 (18.3) | 4 (3.3) | 28 (23.3) |
| 2012 | 42 (40.8) | 20 (19.4) | 3 (2.9) | 19 (18.4) |
| 2013 | 36 (48.6) | 18 (24.3) | 2 (2.7) | 16 (21.6) |
| 2014 | 9 (25.0) | 4 (11.1) | 0 (0.0) | 5 (13.9) |
| 2015 | 3 (42.9) | 2 (28.6) | 0 (0.0) | 1 (14.3) |

Result non-reporting per Year and Study Phase (n=725)
| Study Start Year | Overall, n (%) | Phase 2, n (%) | Phase 2/3, n (%) | Phase 3, n (%) |
| --- | --- | --- | --- | --- |
| 2008 | 72 (50.7) | 49 (34.5) | 6 (4.7) | 17 (12.0) |
| 2009 | 72 (57.1) | 44 (34.9) | 10 (7.9) | 18 (14.3) |
| 2010 | 60 (51.3) | 36 (30.8) | 7 (6.0) | 17 (14.5) |
| 2011 | 66 (55.0) | 48 (40.0) | 6 (5.0) | 12 (10.0) |
| 2012 | 61 (59.2) | 39 (37.9) | 5 (4.9) | 17 (16.5) |
| 2013 | 38 (51.4) | 27 (36.5) | 1 (1.4) | 10 (13.5) |
| 2014 | 27 (75.0) | 18 (50.0) | 3 (8.3) | 6 (16.7) |
| 2015 | 4 (57.1) | 2 (28.6) | 1 (14.3) | 1 (14.3) |

Result reporting per Study Start Year and Sponsor (n=725)
| Study Start Year | Overall, n (%) | Industry, n (%) | Academia/ Non-Profit, n (%) |
| --- | --- | --- | --- |
| 2008 | 70 (49.3) | 49 (34.5) | 21 (14.8) |
| 2009 | 54 (42.9) | 41 (32.5) | 13 (10.3) |
| 2010 | 57 (48.7) | 40 (34.2) | 17 (14.5) |
| 2011 | 54 (45.0) | 43 (35.8) | 11 (9.2) |
| 2012 | 44 (42.7) | 33 (32.0) | 11 (10.7) |
| 2013 | 36 (48.6) | 29 (39.2) | 7 (9.5) |
| 2014 | 9 (25.0) | 8 (22.2) | 1 (2.8) |
| 2015 | 3 (42.9) | 2 (28.6) | 1 (14.3) |

| Study Start Year | Overall, n (%) | Industry, n (%) | Academia/ Non-Profit, n (%) |
| --- | --- | --- | --- |
| 2008 | 72 (50.7) | 33 (22.8) | 39 (28.2) |
| 2009 | 72 (57.1) | 35 (28.0) | 37 (28.8) |
| 2010 | 60 (51.3) | 31 (25.6) | 30 (25.6) |
| 2011 | 66 (55.0) | 28 (23.4) | 38 (32.3) |
| 2012 | 59 (55.0) | 25 (24.5) | 34 (32.1) |
| 2013 | 38 (57.3) | 21 (29.3) | 17 (22.7) |
| 2014 | 27 (75.0) | 14 (38.9) | 13 (36.1) |
| 2015 | 4 (57.1) | 1 (14.3) | 3 (42.9) |

## DECLARATIONS

### Ethics approval and consent to participate

Not applicable.

### Consent for publication

Not applicable.

### Availability of data and materials

The general datasets generated and analyzed during the current study are available in the ClinicalTrials.gov repository, https://clinicaltrials.gov/. Moreover, the specific datasets used and analyzed during the current study are available from Timothy J. Seabrook on reasonable request.

### Competing interests

NKM was employed at F. Hoffmann-La Roche Ltd at the time of study conception, data collection and analysis. JG, RRE, and TJS are employees of and hold shares in F. Hoffmann-La Roche Ltd. We attest that we herein have disclosed any and all financial or other relationships that could be construed as a conflict of interest and that all sources of financial support for this study have been disclosed.

### Funding

This research was supported by F. Hoffmann-La Roche Ltd. The funding body was not part of the study design, data collection, analysis, and interpretation, or in writing the manuscript.

### Authors’ contributions

The study was conceived by JG, RRE, and TJS. Data acquisition, curation and analysis were performed by NKM. The results were interpreted by all authors. The manuscript was compiled by NKM and revised by JG, RRE, and TJS. All authors read and approved the final manuscript.

## Acknowledgements

We would like to thank Dr. Winnie Yeung for advice and assistance with the statistical analysis.

## Footnotes

Not applicable.

## Notes

### Summary of Updates

Figures and tables have been added within the text.

